# Motif2Site: a Bioconductor package to detect accurate transcription factor binding sites from ChIP-seq

**DOI:** 10.1101/2022.09.22.509048

**Authors:** Peyman Zarrineh, Zoulfia Darieva, Nicoletta Bobola

## Abstract

Transcription factors (TFs) binding are in the core of the Regulatory networks studies. ChIP-seq experiments are available for many TFs in various species. As TFs co-bind in cis-regulatory elements regions to control gene expression, studying the existing relation among co-binding TFs such as distance of binding sites or co-occupancy are highly important to understand the regulatory mechanisms. Currently, to detect binding sites of each TF in cis-regulatory elements, first binding regions of each TF are detected by standard peak calling methods, and at the second step the best candidate binding sites are prioritized by motif detection methods in binding regions. However, it is well-known that the best prioritized candidate motifs are not necessarily the actual binding sites of TFs. Furthermore, motif prioritizing methods that consider more genomic features complexities of TFs bindings are usually computationally expensive methods. Here, we tend to improve the TF binding sites accuracy detection by using the original ChIP-seq signal. The motifs which are located closer to the summits of binding region peaks are more likely to be the actual binding sites. Therefore, We developed a novel post-processing Bioconductor package called Motif2Site to detect TFs binding sites from user provided motif sets and recenter them across experiments. We applied Motif2Site method to detect TF binding sites for major mouse embryonic stem cell (mESC) as well as mouse fetal and birth time (P0) heart TFs. Motif2Site could detect binding regions with comparable accuracy to the existing state-of-the-art while it substantially increased the accuracy of the detected binding sites. Motif2Site could future improve the accuracy of binding sites prediction by recentering binding sites across developmental conditions (fetal/P0 heart) and across homologous TFs (ex. GATA4/GATA6 and MEF2A/MEF2C). Purifying high-confidence binding sites in mouse fetal heart, enabled us to study the co-binding properties of TFs in cis-regulatory elements. We could also traced TFs footprints in selected heart-specific VISTA enhancers chromatin accessible regions.

## Introduction

Regulatory networks control gene expression. Transcription factors (TFs) are the backbone of the regulatory networks and control gene expression by binding to DNA. ChIP-seq is the main high-throughput technology to detect the regions that TFs interact with DNA, referred to as binding regions. To detect TF binding regions, short reads of ChIP-seq experiments are aligned to the genome. Then, regions with large number of aligned reads, referred to as peaks, are called with peak calling methods (Thomas et al 2017). The count values in the genomic regions are usually modelled by Poisson or negative binomial distributions (Thomas et al 2017). Model-based Analysis of ChIP-Seq (MACS) is the most used peak calling method (Zhang et al 2008).

Most TFs tend to bind to relatively short DNA sequences referred to as TF binding sites. Studying binding sites of major TFs in cis-regulatory elements (CREs), such as enhancers and promoters, is vital to understand the underlying mechanisms that control gene expression (Wittkopp et al 2012). TFs often co-bind in a specific manner to control the gene expression: order, spacing, orientation, and affinity of binding sites within a CRE are important for TF binding which is referred to as TF binding grammar (Jindal et al 2021). TFs can interact with each other when they co-bind in a close genomic vicinity or compete to their spatially closely located binding sites (Boeva 2016). Furthermore, TF spatial occupancy around its binding sites is another important feature to regulate gene expression. TF Footprinting methods on chromatin accessibility assays such as DNA-seq or ATAC-seq is a major way to study occupancy of TFs (Vierstra et al 2020). A recent publication by ENCODE consortium include TF footprinting of several TFs in different tissues (Vierstra et al 2020). Footprinting is possible at single DNA molecule resolution with a recent developed technology (Sonmezer et al 2021). This makes it possible to study co-occupancy of TFs, and further reveals that whether a pair of TFs compete or collaborate to occupy CREs (Sonmezer et al 2021).

Although detecting accurate TFs binding regions from ChIP-seq is a straightforward task, detecting the actual binding sites of TFs is still a scientific challenge. For this aim, lab-based ChIP-exo approaches such as ChIP-nexus (He et al 2015) method have been developed to detect smaller binding regions, and also TF binding sites detection methods have been improved computationally. TFs generally tend to bind to a set of DNA sequences having high affinity for binding TFs referred to as binding site motifs (Boeva 2016). These TF binding site motifs have been modelled as consensus DNA IUPAC sequence, position weight matrices (PWMs), and k-mers (Boeva 2016). Various methods have been developed to prioritize the best binding sites motifs in detected binding regions from ChIP-seq experiments. HOMER (Heinz et al 2010) and MEME (Machanick) are the most used methods to predict the best binding sites motifs using PWMs. Bayesian networks and other classification methods have also been developed to predict binding sites motifs using k-mer DNA sequences (Boeva 2016). More recently, deep learning methods like BPNet has been developed to improve the accuracy of binding sites motifs prediction (Avsec et al 2021).

Although the computational methods to predict binding sites have been improved substantially, the predicted binding sites by these methods are not necessarily the actual binding sites. It is a well-known fact that TFs are not always bind to the most affine binding site motifs as their co-binding partners play a major role in their binding. Furthermore, ChIP-exo methods have not maturated enough to replace the ChIP-seq experiments. Therefore, in this study we suggest to use the original ChIP-seq signals to prioritize the binding site motifs. Here, the idea is that the actual binding sites are more likely to be located near the summit of the peaks. Therefore, if we perform peak calling on a purified binding sites motifs, it is more likely to detect TF binding sites.

In this study, we introduce an alternative approach to detect TF binding sites by centring peak calling on the user provided binding sites. For this reason, we developed a Bioconductor package called Motif2Site which obtains candidate motifs either as sequence string or bed file from user. Our aim to develop this method is to provide a highly-flexible and user-friendly post-processing method to prioritize high-confidence TF binding sites. We compared the quality of the detected binding regions and binding sites of Motif2Site with **1)** MACS-DiffBind binding region detection followed by HOMER binding site detection and **2)** GEM methods. For this comparison, we used a large set of TFs ChIP-seq experiments in mouse embryonic stem cell (mESC), mouse fetal heart, and mouse birth time (P0) heart systems. We used chromatin accessibility in these system as an independent dataset to verify the binding region quality. While all three methods, MACS-DiffBind, GEM, and Motif2Site predicted binding regions with high accuracy, Motif2Site predicted higher-confidence binding sites overall. Furthermore, the accuracy of Motif2Site predictions was further increased by recentering binding sites across highly similar experiments (like same tissue at fetal and early adult at brith time P0 stages, or ChIP-seq for homologous TFs).

## Materials and Methods

### Motif2Site predicts binding sites and recenter them

Motif2Site performs two major functions: **1)** Detecting binding sites of a TF by performing peak calling around user-provided motif sets by using ChIP-seq (**Figure 1**). **2)** Recentering binding sites across experiments and compare them (**Figure 2**).

**Figure 1.**
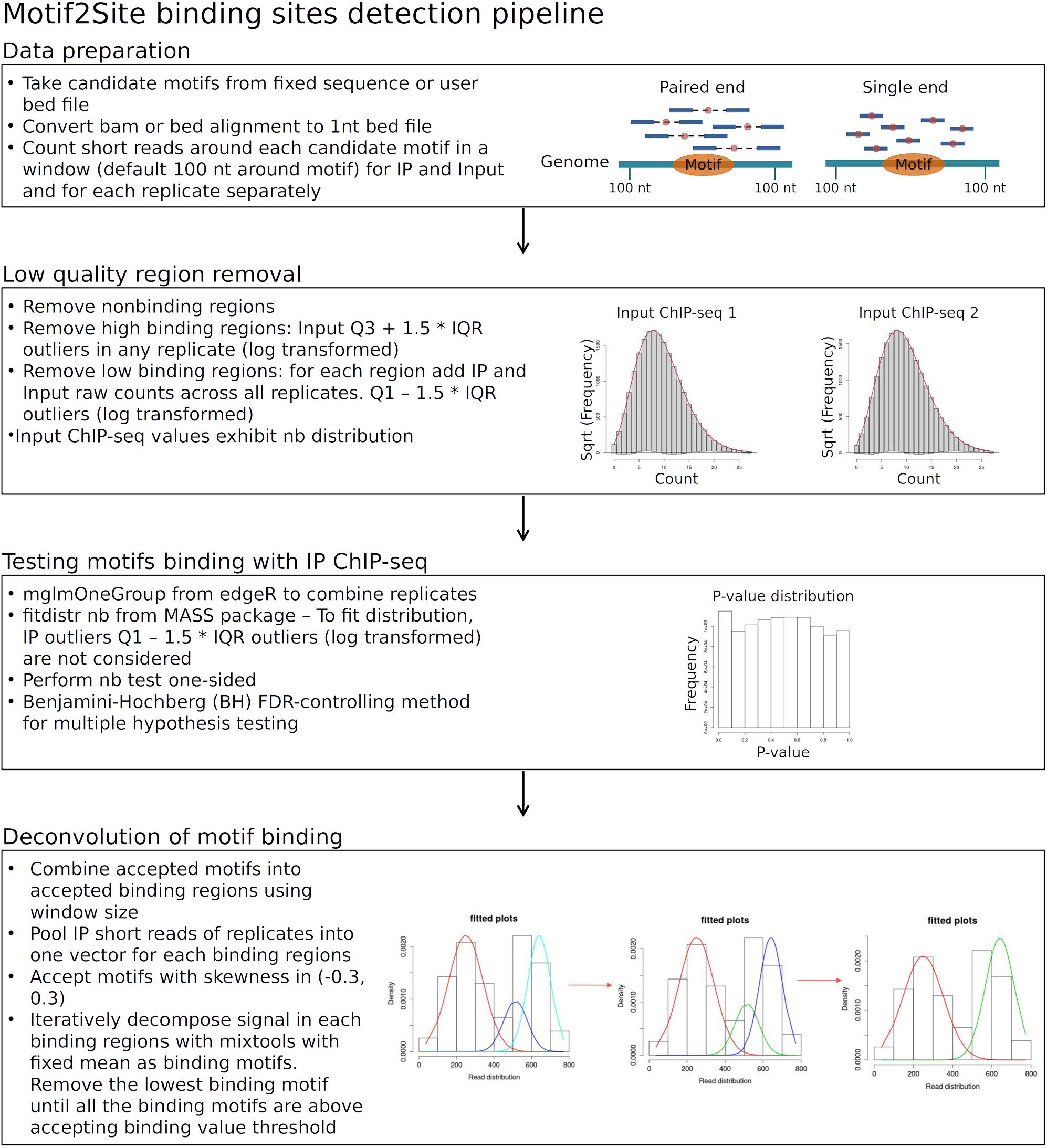
Motif2Site detects binding sites from ChIP-seq data and user provided binding site motifs information. It first preprocesses the input datasets. At the next step, it removes low quality regions using both the IP ChIP-seq and the background (usually Input) ChIP-seq experiments. These low quality regions are detected based on low and high count values, which are more likely to appear due to technical reasons in the ChIP-seq experiments. Then, it performs negative binomial tests to detect candidate binding sites with significant binding intensities. Finally, it deconvolves the adjacent binding site motifs, located in the same binding regions, to verify which of them are the actual binding sites.

**Figure 2.**
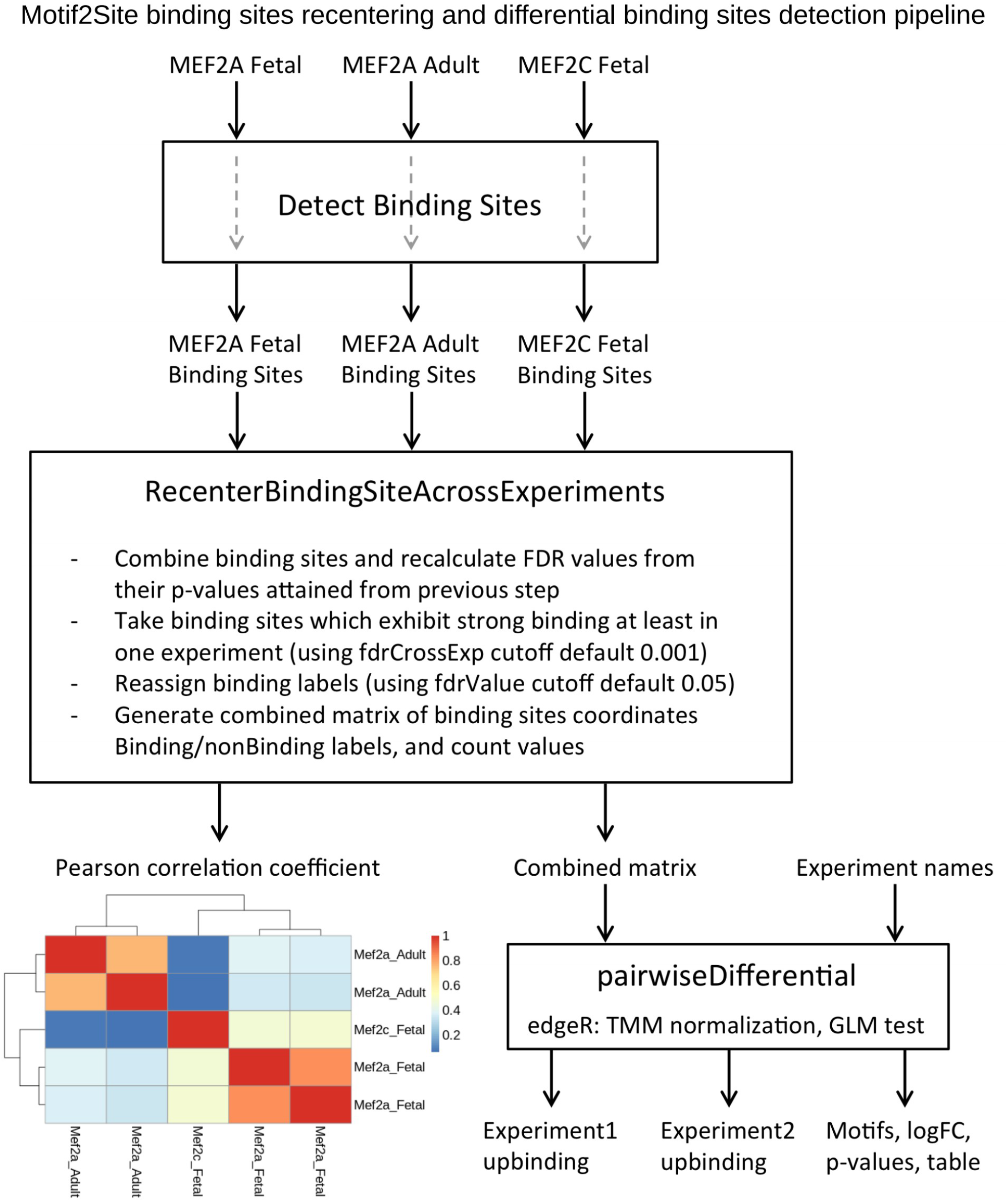
Motif2Site recenters binding sites across conditions, tissues, and homologous transcription factors. It also detects differential binding sites across them by employing edgeR method.

**Figure 1** summarizes the steps of Motif2Site function to call peaks across user provided binding motifs and detect the binding sites. Motif2Site gets two types of input: **1)** ChIP-seq information and **2)** motif information. The first input is ChIP-seq alignment files in BAM or BED format. The second input consists of sequence information either as DNA sequence with mismatch number or a BED file of motifs. This function consists of four major steps: data preparation, low quality regions removal by background ChIP-seq, testing motifs binding with IP ChIP-seq, deconvolution of motif binding. This function generates two output: **1)** files binding sites locations and intensities. **2)** binding regions and the binding sites included in each regions.

In the first step of detecting binding sites function, Motif2Site preprocess both ChIP-seq and binding motif inputs. This function has been implemented in two different forms based on the binding motif input format which can be DNA sequence with mismatch number or a bed file. For the DNA sequence motif input format, Motif2Site finds all the instances of the given DNA sequence motif with user provided mismatch number in the user provided genome using BSgenome package (Pagès H 2022). User can optionally provide regions in a bed format that motifs would only be detected in these regions. This can be open chromatin regions and/or regions with other TFs binding, etc. For the ChIP-seq input files, if the files are in bam format, Motif2Site convert them into bed format by GenomicAlignments package (Lawrence et al 2013). Motif2Site converts alignment information into 1 nucleotide short reads which represent the centers of the reads. Then Motif2Site calculates the binding intensities around motifs for each ChIP-seq separately using a window size (default +/−100 nucleotide around motif).

In the next step of detecting binding sites function, Motif2Site filters non-sequenced, lowly-sequenced, and highly-sequenced regions using count distribution in both IP and background (Input) datasets. First, all the motifs with no binding in both IP and background (Input) files in all replicates of experiments would be removed as nonbinding regions. After removing zeros, background (Input) ChIP-seq experiments, are used to remove the regions which gain high binding intensities due to genomic features. For each background (Input) ChIP-seq, high-intensity binding outliers were detected as high binding regions (using Q3 + 1.5 * IQR). After removing high binding regions, Motif2Site removes the low binding regions due to genomic features across all experiments. Motif2Sites pools all the binding values across IP and background (Input) replicates for each motif. Then it detects low binding outliers at log-scale (using Q1 - 1.5 * IQR). Motif2Site reports the number of motifs filtered as nonbinding, high binding, and low binding in its output folder. After filtering the ChIP-seq experiments exhibit negative binomial distribution.

For the motifs, located in the same binding regions, the binding signals are deconvolved to verify which of them are the actual binding sites as the final step of detecting binding sites function. For this aim, first the binding short reads of different IP ChIP-seq experiments around accepted binding regions are pooled together. Then, these binding signals are deconvolved across motifs, located in the same binding regions, to the Gaussian peaks by using mixtools package in R (Benaglia et al 2009). Motifs are located as the center of Gaussian peaks. If the motifs with lowest binding signal after decomposition exhibit low binding affinity, Motif2Site removes this motif and repeat the deconvolution process. Here, the lowest intensity value across all accepted binding motifs is used as the low binding affinity cutoff. As the signal deconvolution is a computational expensive process, Motif2Site removes motifs with skewed binding distribution around them prior to deconvolution. For this aim, Motif2Site uses third standard moment with 0.3 as cutoff. Motif2Site performs this iterative procedure for all binding regions which include more than one binding site. Motif2Site reports the number of binding motifs which are filtered based on skewness and the Gaussian decomposition as part of its output.

Motif2Site also recenter peaks across tissues, condition, and TF homologs **Figure 2**. This function accept the results of binding site detection function and combine them across experiments. Motif2Sites pools binding sites across experiments which exhibit high-confidence binding at least in one experiment by using a stringent user-defined FDR cut-off (default 0.001). Then it considers the p-values of these binding sites for each experiment separately, and it performs BH correction once again for the pooled binding sites for each experiment separately. For each experiment, Motif2Site accepts binding sites by using another user provided FDR threshold (default 0.05). Motif2Site summarizes the binding information across different experiments in a combined matrix. Motif2Site also calculates pairwise Pearson correlations across experiments using count values at log-scale. Motif2Site also facilitate differential binding study of pairs of experiments across recentered experiments. It fetches the raw count values of binding intensities of two experiments, which should be compared, from the combined matrix of the recentered experiments. Differential binding sites across two user provided conditions are computed by employing edgeR (Robinson et al 2010) TMM normalization followed by glmLRT test (Robinson et al 2010).

### Processing ChIP-seq and ATAC-seq datasets

mESC and heart fetal and birth time (P0) ChIP-seq and ATAC-seq accession numbers are summarized in **Table S1**. mESC SOX2, OCT2, and NANOG ChIP-seq and ChIP-nexus have been published in (Avsec et al 2021). Bam alignment and peaks of mESC ChIP-seq as well as summits of ChIP-nexus peaks were downloaded directly from GEO database. Similarly, ATAC-seq regions of mESC published in (Friman et al 2019) were downloaded from GEO database. “GSE134652_ZHBTc4_2TS22C_mergepeaks.bed” file was used as accessible crhomatin regions in mESC system.

We have used a comprehensive ChIP-seq data in mouse fetal and at birth (P0) heart including MEF2A, MEF2C, TEAD1, SRF, TBX5, NKX2.5 ChIP-seq data from Akerberg et al 2019. In addition, we added GATA4 mouse fetal and P0 heart (He et al 2014), TBX20 fetal heart (Boogerd et al), TBX20 adult heart (Shen etl al 2011), GATA6 posterior branchial arches (PBA) (Losa et al 2017), and CTCF heart at birth (P0) from ENCODE project (ENCODE Project Consortium). To analysis heart ChIP-seq experiments, the short reads were aligned to mouse genome mm10 using Bowtie2 (Langmead et al 2012). Samtools (Li et al 2009) was used to remove the aligned reads with a mapping quality lower than Q10. ATAC-seq of heart fetal (Akerberg et al) and P0 from ENCODE (ENCODE Project Consortium) were analyzed by esATAC (Wei et al 2018) and recentered by DiffBind (Ross-Innes et al 2012) Bioconductor packages.

TF ChIP-seq experiments were analysed by Motif2Site and compared with MACS-DiffBind-HOMER pipeline and GEM. Motif2Site was used both by string motifs shown in **Table S2** and motif bed files generated by HOMER method (Heinz et al 2010) using motifs and LOD values summarized in **Table S3**. Narrow peaks called by MACS method for all the TF ChIP-seq. Heart TFs and mESC TFs are recentered in two separate table by DiffBind method (Ross-Innes et al 2012). For each TF, HOMER (Heinz et al 2010) was used to detect best binding sites in MACS-DiffBind binding regions. For this aim, we used HOMER motifs in **Table S3** without using the fixed LOD values. In each region only one motif (the best motif), with the highest LOD value, was reported. GEM (Guo et al 2012) was run by default settings.

To perform comparative analyses, we intersected TF binding regions and binding sites across methods using ChIPpeakAnno Biocondcutor package (Zhu et al 2010), and visualized the intersections as Venn diagrams using nVennR package (Quesada 2021). We also intersected each TFs binding with the relevant ATAC-seq data using Bioconductor GenomicRanges package (Lawrence et al 2013). The distance between binding sites were also calculated using GenomicRanges package (Lawrence et al 2013). ggplot2 (Wickham 2016) was used to generate all comparative plots, and pheatmat package (Kolde 2019) was used to generate TFs pairwise distance heatmap.

H3K27ac marker from (Nord et al 2013) datasets were used, to compare activity across heart, forebrain, and liver at mouse developmental stages. Similar to TF ChIP-seq, the short reads were aligned to mouse genome mm10 using Bowtie2 (Langmead et al 2012), and Samtools (Li et al 2009) was used to remove the aligned reads with a mapping quality lower than Q10. Broad peaks called by MACS method for all the histone ChIP-seq, and the regions were recentered by DiffBind method (Ross-Innes et al 2012). RPKM values were used to calculate the fold changes across tissues for each region in each developmental time. For the replicated experiments, average RPKM values were used for fold change calculations. pheatmat package (Kolde 2019) was used to plot log fold change patterns of heart vs liver and forebrain through the temporal development for the desired active regions.

## Results

### Motif2Site predicts binding regions and binding sites with high accuracy

We have developed Motif2Site, a novel Bioconductor package, to select binding sites from user provided TF binding motifs and recenter them across experiments (**Figure 1** and **Figure 2)**. To detect binding sites, this method takes user provided input TF binding motifs information either as DNA string with mismatch number or as a bed file. To facilitate choosing TF binding motifs by the user, Motif2Site provides a function to compare the motif regions with high-confidence user provided binding regions. In this study, we considered TFs MACS-DiffBind binding regions intersected by open chromatin accessible regions as high-confidence binding regions. Two DNA string sequence examples with mismatch number sets are highlighted in **Figure S1A** to choose which motif to be used SOX2 and OCT4. In these plots, The Y-axis show how much of high-confidence binding regions (MACS-DiffBind intersected by ATAC) are covered by this motif. X-axis show the fraction of binding site motifs covered by high-confidence binding sites. User can choose the desired binding sites motifs in terms of precision and recall using these plots. After running Motif2Site, this method generates aggregated binding short reads distribution around binding sites which are far from other binding sites and their signals do not need to be deconvolved. This aggregated short read distribution can also be used as another measure that can assists users to decide about motifs or parameter setting values. **Figure S1B** shows this binding distribution around OCT4 binding sites using FDR values equal to 0.05 and 0.005 for the used string motif. As it can be seen in **Figure S1B**, the default value 0.05 does not detect binding sites with good short read signal around them. For this reason we used FDR 0.005 for OCT4 binding site detection. OCT4 was an extreme example in this study for which purifying binding site was hard for both sequence and HOMER motif inputs, and for the rest of TFs we used motifs (**Table S2** and **Table S3**) with default FDR value 0.05.

For all TFs in this study, Motif2Site, MACS-DiffBind, and GEM could predict high-confidence binding regions for all systems. First of all, the binding regions that these three methods detected were highly similar. **Figure 3A** shows the similarity between common regions detected by Motif2Site DNA string input, Motif2Site HOMER motif bed input, MACS-DiffBind, and GEM for mESC TFs. 7,962 SOX2, 18,852 OCT4, and 30,536 NANOG binding regions were detected by all methods. On the other hand, binding regions numbers detected only by one method were much smaller than the binding regions detected by more than one method (**Table S4** mESC TFs). Binding regions were also compared to ATAC-seq open chromatin regions as independent datasets to further verify the accuracy of predictions (**Figure 3B**). ATAC-seq verified that the majority of mESC TF binding regions detected by all the methods are accessible chromatin. Also it shows that the methods exhibit high confidence and comparable results in terms of precision and recall. The same was also true for mouse fetal and P0 heart TFs binding sites (**Figure S2** and **Figure S3**). **Table S4** summarises the ratio of detected binding regions by different methods in both mESC and heart systems. It shows that the majority of binding regions have been detected by more than one method for different TFs which is another indication of high-confidence binding regions detection by different methods. Furthermore, for almost all the TFs in this study, the dominant majorities of detected binding regions by any of the methods have been detected by at least another method (**Table S5**). Verification of these high-confidence binding regions, derived by each method, with ATAC-seq regions exhibits comparable results in terms of precision and recall for mouse heart fetal and P0 TFs (**Figure S2**).

**Figure 3.**
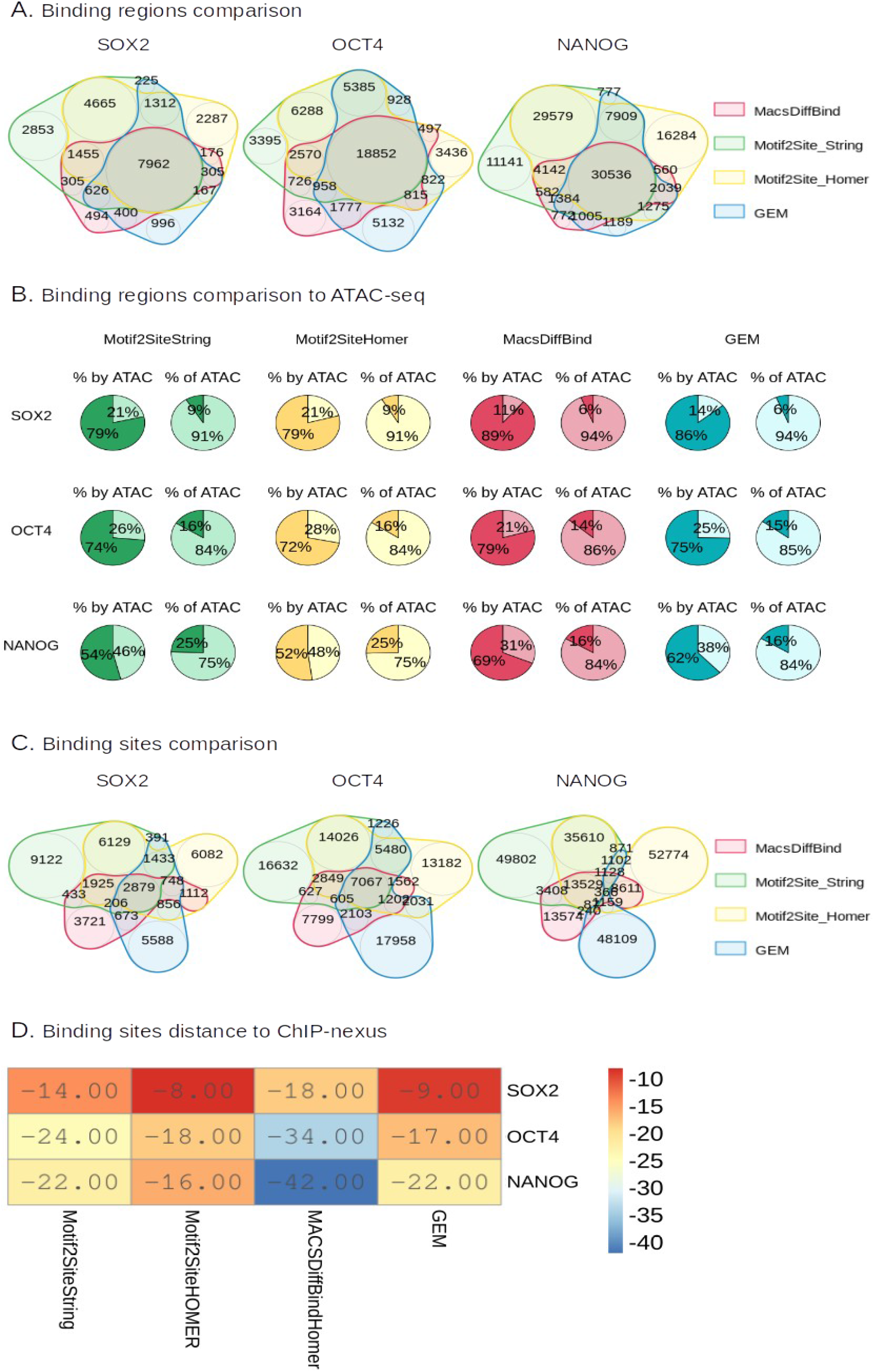
Binding regions and binding site comparisons across methods for SOX2, OCT4, and NANOG in mESC. **A.** Binding region comparison across methods **B.** Accuracy comparison by comparing detected binding regions by each method to accessible chromatin from ATAC-seq **C.** Binding sites comparison across methods. **D.** The median distance of binding sites, detected by different methods, to binding summits of ChIP-nexus experiments.

Although different methods predict high confidence and highly similar binding regions, they predict largely different binding sites. For SOX2, OCT4, and NANOG in mESC system, the number of binding sites detected by more than one method is smaller than the number of binding sites detected by only one method individually (**Figure 3C** and **Table S6** mESC TFs). However, still the majority of binding sites detected by each method were detected by at least another method (**Figure 3C** and **Table S7** mESC TFs). Here the only exception is GEM NANOG binding sites, and in general GEM detected more unique binding sites than binding sites also detected by other methods (**Figure 3C** and **Table S7** mESC TFs). On the other hand, Motif2Site with HOMER motif bed input gained the highest intersecting binding sites (**Figure 3C** and **Table S7** mESC TFs). Similarly, detected binding sites were highly different across different methods in mouse fetal and P0 heart (**Figure S3** and **Table S6**). However similar to mESC, a considerable ratio of detected binding sites, detected by one method, have been detected by at least one another method (**Table S7**). Motif2Site DNA string input and Motif2Site HOMER motif bed input gained the highest ratio of reproducible binding sites by at least another method (**Table S7**). Here, we could expect that Motif2Site from two different sets of binding site motifs predict non-similar binding sites as the initial binding site motifs are far from each other. However, Motif2Site could predict more similar binding sites as it could rejects non-binding sites from initial binding site motifs sets.

As the ChIP-nexus binding summits are available for the major mESC TFs, we compared the binding sites predicted by all methods to these summits. To this aim, we used regions detected as binding regions by all methods and included binding summits from ChIP-nexus datasets. **Figure 3D** exhibits the median distance of the center of binding sites from different methods to these summits in the mentioned regions. Motif2Site with HOMER motif detected the closest binding sites to the ChIP-nexus. GEM exhibited the second-best performance while the best HOMER motifs in MACS-DiffBind regions exhibited the worst performance. This clearly shows that the best HOMER motif in the binding regions are not necessarily the binding sites, at least in these examples.

Motif2Site recenters binding sites across experiments (**Figure 2**), and recentering binding sites across experiments further increase the accuracy of the predicted binding sites. **Figure S4** shows the increased accuracy of binding regions after recentering in mouse heart TFs. More than 70% of all detected regions by Motif2Site method intersect with ATAC-seq regions (precision more than 0.7) after recentering for both DNA string and HOMER bed file motif inputs. Among TFs examined, SRF was the only exception which retains precision less than 0.7. We realized that the MAD box which is the best enriched motif in SRF binding regions, **Figure 4A** top panel, is not the actual binding site of the ChIP-seq experiments, and an ETS-like motif is the actual binding sites of the used ChIP-seq experiments (**Figure 4A** lower panel). By looking at the binding site signal distribution around the MAD box motifs which did not need to be deconvolved (**Figure 4B** left panel), we observed that the real binding sites are located in average around 27 nucleotides (nt) distant to the ChIP-seq binding sites. For this reason, we took DNA nucleotide sequences with 27nt distance to the called binding sites by Motif2Site HOMER bed input, and run a fresh HOMER de novo motif search on them to detect the actual binding sites of SRF ChIP-seq. Consistent with cooperation between SRF and ETS (Hipskind et al 91), we identified ETS-like motif in **Figure 4A** lower panel. **Figure 4B** right panel shows that this motif is indeed provides the correct binding sites for SRF ChIP-seq, and the binding signals around the non-deconvolved motifs in Mtoif2Site Bed HOMER input results exhibit a sharp peak distribution around these binding sites. **Figure 4C** shows that using ETS, rather than SRF motif, increases the precision of binding regions compared to ATAC-seq over 0.7 for both fetal and birth time P0 adult stages. We run HOMER to find de novo motif on new SRF fetal binding sites obtained by Motif2Site HOMER bed input. **Figure 4D** shows that the enriched motif was even further improved, and this time it resembles of a combined ETS-BHLH motif.

**Figure 4.**
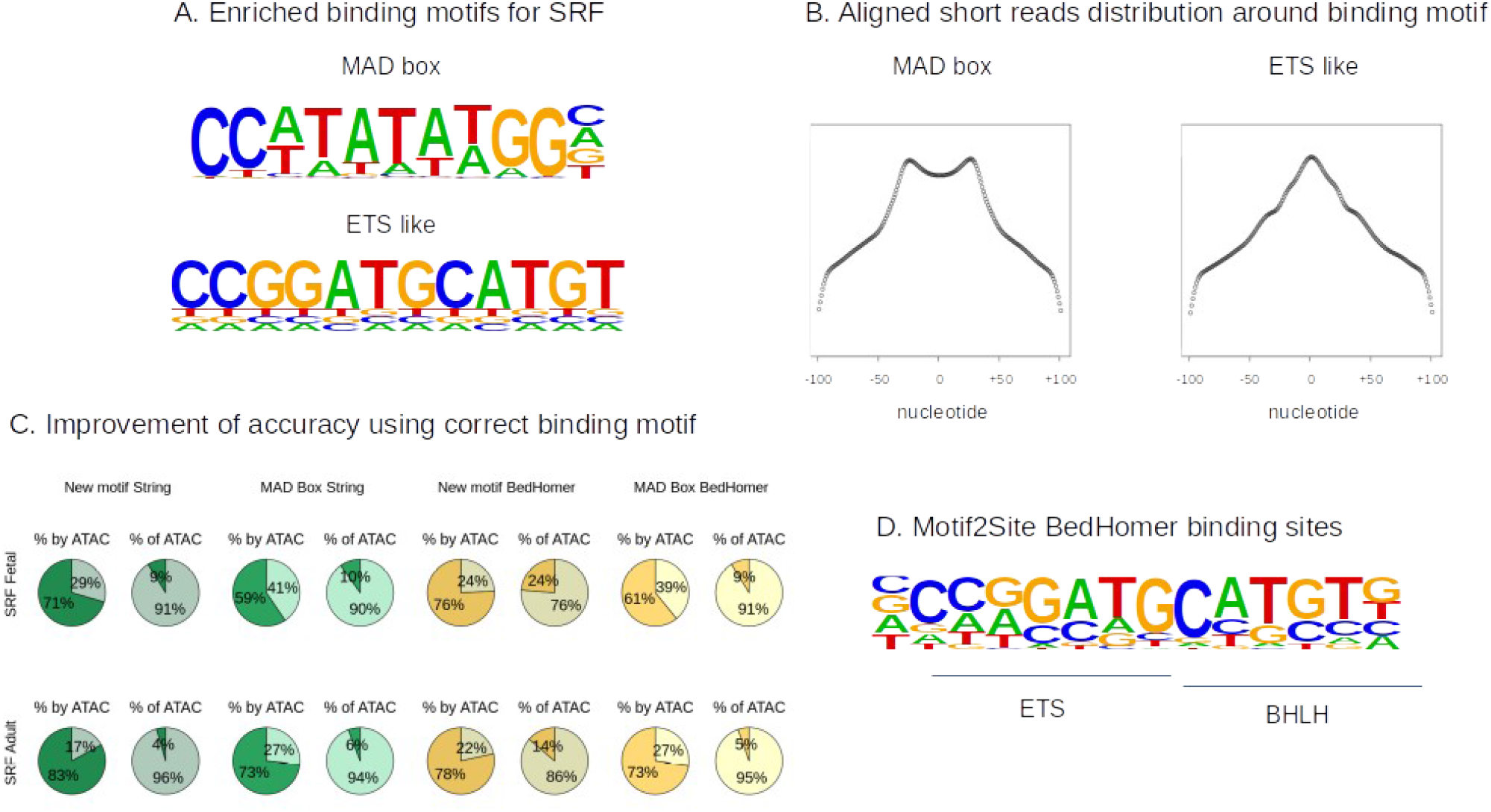
A novel SRF binding sites motif. **A.** Well-known MAD-box binding sites motifs is highly enriched in SRF binding regions (top panel) while an ETS-like motif exist appear roughly around 27 nucleotide from this motif. **B.** Binding site signal distribution around the MAD box motifs (left panel) and novel ETS-like motif (right panel). The binding signal is clearly centered around ETS-like motif. **C.** Motif2Site detected SRF binding regions comparison using MAD-box and ETS-like motifs to open chromatin regions from ATAC-seq. Motif2Site binding regions detected by using ETS-like motif has higher accuracy. **D.** Running HOMER on Motif2Site binding sites using bed HOMER input resulted to a more accurate novel motif which contains ETS and BHLH domains.

### Motif2Site reveals co-binding modules in mouse developing heart

As Motif2Site provides high-confidence binding sites in heart (precision larger than 0.7), we combined Motif2Site results from string and HOMER bed file inputs to study TFs co-binding in accessible chromatin and its effect on heart-specific enhancers activities. **Figure 5A** shows combinations of TFs co-binding in ATAC-seq regions in fetal mouse heart: the majority of accessible chromatin regions are targeted by combinations of TFs. To analyse the contribution of TF combinations to CRE activity, we calculated the average log-fold change (logFC) of H3K27ac (a mark of active enhancers) in mouse fetal heart versus forebrain and liver. We observed 23 TF combinations associated with higher H3K27ac values in heart relative to forebrain and liver at all developmental stages. **Figure S5** shows the average logFC values for these combinations through the developmental stages. These 23 TF combinations appear in a total of 5637 CREs (**TABLE S8**). Higher logFC of these TF combinations in heart versus two other tissues at all developmental stages suggests that these heart-specific TF co-bindings increase enhancer activities in mouse fetal heart and consequently controls gene expression in a tissue-specific manner.

**Figure 5.**
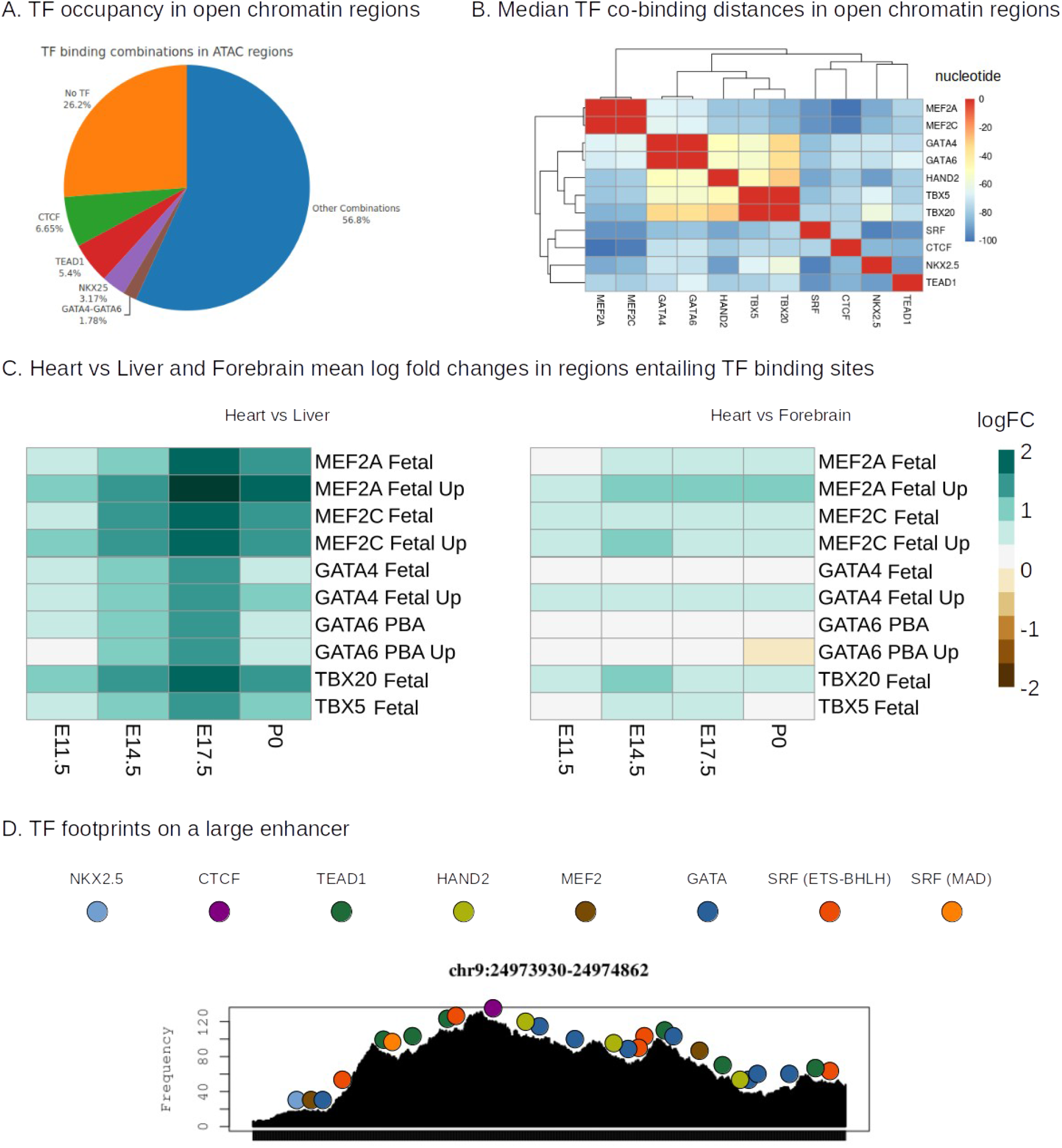
TF cobinding properties in mouse fetal heart. **A.** TF combinatorial binding ratio in open chromatin regions. The majority of accessible regions are bound by combinations of heart TFs. **B.** Median distances between closest pairs of TFs in open chromatin regions. **C.** H3K27ac log-fold changes in heart versus liver (top panel) and heart versus forebrain (lower panel) throughout the developmental stages. The log-fold changes were calculated for regions containing binding sites or differential binding sites of given TFs, as identified by separate colours. **D.** TF footprints on a large heart-specific open chromatin which consists of two intersecting VISTA enhancers that control TBX20.

Motif2Site predicted binding sites also facilitated the study of TFs co-binding and occupancy in mouse fetal heart system. First for each pair of TFs, we calculated the median distance of their co-binding in chromatin accessible regions. To this aim, we consider pairs of binding sites, which are closest to each other in ATAC-seq regions. In other worlds, binding site B1 of TF1 and binding site B2 of TF2 are considered as paired if they are located in ATAC-seq regions (extended 100 nt from each side to be more inclusive), and B2 is the closest binding site of TF2 to B1, and B1 is the closest binding site of TF1 to B2 (**Figure S6**). **Figure 5B** summarizes the median binding distances of TFs in ATAC-seq regions in a heatmap. Generally we did not observe a fixed distance of binding events between two pairs of TFs. However, we found that many TF binding sites are located in close vicinity. This is especially true for GATA, TBX, and HAND2 binding sites which are known to interact with each other (Delgado et al 2021, Losa et al 2017).

Differential binding intensities across homologous TFs is another approach to study TFs co-binding. Motif2Site provides binding intensities correlation and differential binding across homologous TFs as well as across developmental stages in mouse heart. **Figure S7A** shows that MEF2A and MEF2C binding intensity values exhibit medium correlation in mouse fetal heart while MEF2C fetal binding intensity values do not exhibit any correlation with MEF2A adult (at birth time P0). This indicates that MEF2 binding is more related to the developmental stage than which homologous are binding to the binding sites. In a similar way, GATA4 fetal heart binding intensity values exhibit low correlation with GATA6 PBA, while GATA4 adult (at birth time P0) binding intensity values do not exhibit any correlation with GATA6 PBA. In this case PBA and fetal heart are related but not identical tissues. However, GATA4 fetal binding intensities show low correlation with both GATA4 P0 and GATA6 PBA binding intensities. This shows that the developmental stage, type of tissues, and TF homologous have effect on the binding sites of GATA. We used Motif2Site to detect differential binding sites across GATA4 fetal heart and GATA6 PBA as well as MEF2A and MEF2C fetal heart. We run HOMER (+/− 50 nt) around differential binding sites of related TFs (summarised in **Figure S7B**). The enriched motifs in differential GATA binding (GATA4 vs GATA6) were tissuespecific motifs. In particular, GATA4 fetal heart is associated with its heart-specific co-binding partners. Interestingly we observed that MEF2C display higher binding levels with heart-specific co-binding partner TFs (**Figure S7B**). In contrast, MEF2A binds with higher intensities when cobinds with bzip factors (both AP1-like and CREB/ATF families like) (**Figure S7B**). As Motif2Site provided us with binding sites of different heart TFs, we counted binding sites percentage of different TFs in GATA and MEF2 differential regions (**TABLE S9**). The ratio of observed co-bindings further verifies the observed difference of GATA4 fetal heart and GATA6 PBA as well as MEF2A and MEF2C fetal heart in motif search. By using H3K27ac logFC of heart versus forebrain and liver, we observed that differential MEF2C and MEF2A exhibit more activity than non differential MEF2 binding sites (**Figure 5C**). Similarly, TBX20 binding sites exhibit stronger activity in heart than TBX5 (**Figure 5C**). We did not perform TBX5 and TBX20 differential binding analysis because TBX20 binds to a small portion of TBX5 binding sites (TBX20 detects a longer and more specific binding site than TBX5 one). Although GATA binding in general does not increase heart enhancer activity as much as MEF2 and TBX binding (**Figure 5C**), but fetal heart is still more active in the regions with binding intensities GATA4 fetal > GATA6 PBA (from Motif2Site differential) than other GATA binding regions (**Figure 5C**).

As mentioned above, purifying high-confidence TF binding sites would facilitate the study of TF occupancy. Therefore, we traced the occupancy of heart-specific TFs provided by Motif2Sites in heart-specific enhancers to demonstrate how high-confidence TF binding sites provided by Motif2Site can facilitate the TF occupancy studies. For this reason, we considered active VISTA enhancers (Visel et al 2007) in developing mouse around TBX20 to perform footrpinting. Four VISTA enhancers, with H3K27ac logFC larger than 1 in heart compare to both forebrain and liver for all fetal developmental stages, are located around the TBX20 locus. To visualize the footprints on chromatin accessible regions, we pooled two fetal mouse heart ATAC-seq. Two out of the four VISTA enhancers are intersecting and they include a highly accessible regions with large number of TF binding events (**Figure 5D**). We can observe some long footprints (in terms of DNA nucleotide numbers) due to co-binding of several TFs in this example (**Figure 5D**). **Figure S8** include all four VISTA enhancers (two of them intersecting in their genomic coordinates) and TF binding sites. In these examples, GATA binding and HAND2-GATA co-binding exhibit clear footprints. Interestingly we observe clear footprints for many detected TF binding sites detected by Motif2Sites. The length and depth of footprints are highly varied especially for tandom co-binding of TFs. This shows the footprints in open chromatin and TF binding sites are both are needed to be considered together for a comprehensive occupancy study which can be facilitated by a high-confidence binding sites detections method like Motif2Site method.

## Discussion

In this paper, we introduced Motif2Site as a new post-processing Bioconductor package to detect TF binding sites from ChIP-seq experiments by using user provided motifs. We used two data-rich systems, mESC and mouse heart and compared Motif2Site performance with MACS-DiffBind and GEM methods. Using ATAC-seq as an independent dataset to verify detected binding regions, Motif2Site achieved comparable results in terms of precision and recall to the other methods. The resolution of results was especially high for the TFs which bind to a clear and specific binding sites in their binding regions. Motif2Site outperform MACS-DiffBind-Homer and GEM in terms of binding site detection as its detected binding sites are more reproducible when used with two different input sets String motifs and HOMER motifs. Also, Motif2Site in combination with HOMER motifs predicts the closest binding sites to the high-confidence binding summits of ChIP-nexus in mESC system.

Motif2Site has been developed to increase the accuracy of detecting TF binding sites. The main focus of developing this method was to provide a highly flexible post-processing method for the users to study the binding and co-binding of TFs. As such, Motif2Site provides opportunities to study the grammar of binding and co-binding of TFs. Indeed, in this paper, we studied the binding grammar of the main TFs active in mouse fetal heart. We observed these TFs co-bind in the majority of heart enhancers. While the distance between TF binding sites varies, the median distance of TFs known to co-bind, tend to be shorter than the other combinations. This include the co-binding network of GATA, TBX, and HAND2 factors, which is likely to be orchestrated by MEIS (Delgado et al 2021, Losa et al 2017). We also observed co-binding of SRF with BHLH and ETS factors based on the close vicinity of a MAD box to the major SRF binding sites (Wenwu et al 2011). It is well-known that SRF interacts and co-bind with ETS factors especially ELK1 (Branko et al 1996, Odrowaz et al 2012) and BHLH factors (MOLKENTIN etl al 1996). However, we found a novel ETS-BHLH motifs which appear near the well-established SRF-MAD box combination. Interestingly, this novel motif was the one that was captured by the used ChIP-seq experiment (Akerberg et al 2019).

Purified binding sites with binding intensities have further usages such as deciphering the effect of homologous TFs binding in a system and studying TFs footprints. By using Motif2Site recentering and differential binding functions, we observed that MEF2C co-binds with the main heart TFs with stronger intensity than MEF2A. In contrast, MEF2A co-binds with bzip TFs with higher binding intensities than MEF2C. Having differential binding sites rather than differential binding regions (from a method like MACS-DiffBind) have this advantages that we could perform the motif search around the binding sites with smaller window size. For example, a preferred window size of motif detection in differential binding regions is 200 nucleotide while we used 100 nucleotide window size. We could even make the window size smaller as we had the binding sites of the TFs of interest in the center of regions used for motif search (example +/− 30 nucleotide around binding sites). Having binding sites also increase the resolution of footprinting studies. We looked at the footprint of four VISTA heart specific enhancers in this study using the available bulk ATAC-seq. We observed large footprints in size and depth due to the co-binding of major heart TFs binding in these enhancers. This clearly shows having high-confidence binding sites from a method like Motif2Site would indeed improve the footprinting studies. This framework can also be expanded using single-cell technologies like snATAC-seq to obtain footrpinting resolution at cell-type level.

In summary, Motif2Site provides flexibility for post-processing analysis of regulatory network studies. We have used this method for ChIP-seq datasets of mouse mESC, fetal heart, and P0 heart systems. Importantly, it can also be used for similar technologies to ChIP-seq with an adapted binding input. Similarly, the binding site motif input for Motif2Site method can also be improved by the users. For example, users can provide motif input obtained with state-of-the-art-methods like deep learning models (Avsec et al 2021) or combine motif sets obtained with more than one method. Motif2Site method can employ binding sites from different input sets or across experiments and tissues; in fact, recentering across experiments would even increase the accuracy of detected binding sites. This makes this method especially suitable to build compendia of transcription factor binding sites for TF with several datasets.

## Data and code availability

Source codes, datasets, and intermediate results are available at https://figshare.com/articles/dataset/Motif2Site_paper_support/20735977

## Acknowledgements

This research was supported by Biotechnology and Biological Sciences Research Council (BBSRC) grant BB/T007761/1. The authors would like to thank Prof. Magnus Rattray for the guidance and advice that he provided. The authors would like to thank Dr. Zhabiz Vilkiji for providing codes for visualization and helping to generate figures.

## Supplementary Figures captions

**Figure S1.** Binding site motif selection for Motif2Site. **A.** Comparison of motifs regions to high-confidence binding regions generated by Motif2Site. DNA String motifs for SOX2 and OCT4 were compared to MACS-DiffBind binding regions of SOX2 and OCT4 in ATAC-seq chromatin accessible regions. X-axis shows the ratio of motif sites covered by high-confidence binding regions, and Y-axis shows the ratio of high-confidence binding regions covered by motifs. 0.95 value in Y-axis is highlighted by a red dashed line. **B.** Aggregated binding short reads distribution around the detected OCT4 binding sites from DNA string input which are far from other binding sites and whose signals do not need to be deconvolved. A better signal distribution can be observed for binding sites which were derived with more stringent cut-off (FDR 0.005).

**Figure S2.** Accuracy comparison by comparing to accessible chromatin from ATAC-seq with binding regions in mouse fetal and early adult birth time P0 heart TFs detected by different methods.

**Figure S3.** Binding regions and binding sites comparison in mouse fetal and early adult birth time P0 heart TFs detected by different methods.

**Figure S4.** Accuracy comparison by comparing to accessible chromatin from ATAC-seq of Motif2Site detected binding regions before and after recentering across fetal and early adult birth time P0 stages as well as homologous TFs. Recentering clearly increases accuracy in terms of precision (% by ATAC) and recall (% of ATAC).

**Figure S5.** TF combinations for which the average log-fold change values (Y-axis) were higher in heart than forebrain and liver in all developmental stages (X-axis).

**Figure S6.** A schematic representation of how to pick the pairs of TF co-bindings in an open chromatin region.

**Figure S7.** Motif2Site recentering and differential binding detection from bed HOMER motifs input. **A.** Pearson correlation coefficient calculated by Motif2Site on log-count values for recentered GATA6 PBA, GATA4 fetal, GATA4 early adult birth time P0 (right panel) and MEF2C fetal, MEF2A fetal, MEF2A early adult birth time P0 (left panel). **B.** Enriched HOMER de novo motifs around differential binding sits of GATA6 PBA vs GATA4 adult and MEF2C fetal vs MEF2A fetal using all recenters GATA and MEF2 binding sites as background of de novo motif searches, respectively.

**Figure S8.** TF footprints on four heart-specific VISTA enhancers interacting with TBX20. The lowest panel shows the four heart-sepecific VISTA enhancers locations and their coordinates. The other panels show the ATAC-seq mouse heart fetal cutsites (replicate1 and replicate 2 were pooled). Two of the enhancers intersect and the whole combined region is a large open chromatin for this reason. Therefore, we combined them into one panel. TFs were shown over their binding sites detected by Motif2Site.

## Supplementary Tables captions

**Table S1.** Accession numbers of all ChIP-seq, ATAC-seq, and ChIP-nexus experiments used in this paper.

**Table S2.** DNA strings and mismatch numbers used as input for Motif2Site.

**Table S3.** HOMER motifs with used LOD values to generate bed files as input for Motif2Site.

**Table S4.** The ratio of detected binding regions by different methods in mESC as well as fetal and P0 mouse heart systems.

**Table S5.** the ratio of detected binding regions by each method that have also been detected by at least another method.

**Table S6.** The ratio of detected binding sites by different methods in mESC as well as fetal and P0 mouse heart systems.

**Table S7.** the ratio of detected binding sites by each method that have also been detected by at least another method.

**Table S8.** TF combinations for which the average log-fold change values were higher in heart than forebrain and liver in all developmental stages. The second column shows the number of open chromatin regions in mouse fetal ATAC-seq which bound by these TF combinations.

**Table S9.** The ratio of binding regions in X-Axis entail TFs binding sites in Y-axis. The difference in the rario’s would highlight the difference of co-binding in regions that MEF2A and MEF2C in fetal heart as well as GATA6 PBA and GATA4 fetal hear bind with differential intensities.

